# Antibiotic Use and Resistance in Zambia’s Aquaculture Sector: Assessing Knowledge, Attitudes, and Practices Among Fish Farmers

**DOI:** 10.1101/2025.04.07.647520

**Authors:** Kunda Ndashe, Geoffrey Mainda, Chitwambi Makungu, Mark Caudell, Katendi Changula, Niwael Mtui-Malamsha, Tabitha Kimani, John Bwalya Muma, Masuzyo Nyirenda, Mwaka Sinkala, Bernard Mudenda Hang’ombe

## Abstract

Aquaculture in Zambia is rapidly growing, contributing significantly to food security and income generation. However, the intensification of fish farming raises concerns about antimicrobial use (AMU) and the emergence of antimicrobial resistance (AMR). This study assessed the knowledge, attitudes, and practices (KAP) of fish farmers regarding AMU and AMR across 25 districts in Zambia’s ten provinces. Data were collected from fish farmers using a structured questionnaire distributed via district fisheries officers and analyzed through descriptive and inferential statistics, including logistic regression.

The results revealed significant gaps in knowledge, with only 17.4% of farmers classified as knowledgeable about AMU and AMR. Attitude assessments showed that 60.5% exhibited positive attitudes toward responsible AMU practices, while the remaining 39.5% displayed negative perceptions. In terms of practices, 76.5% adhered to good practices, including consulting veterinary professionals and responsible antibiotic use, whereas 23.5% engaged in poor practices. Key factors influencing KAP outcomes included age, farming experience, and annual production capacity. Farmers aged 30–39 years and those with 1–5 years of experience demonstrated significantly more positive attitudes (p < 0.05), while those with higher production capacities (501–1000 kg) exhibited better practices (p < 0.001).

The findings highlight critical knowledge gaps and inconsistent practices regarding AMU and AMR among Zambian fish farmers. Targeted interventions, including education programs, enhanced access to veterinary services, and strengthened regulatory frameworks, are essential to promote responsible AMU, mitigate AMR risks, and ensure the sustainability of aquaculture in Zambia.

## 1. Introduction

Global aquaculture has undergone remarkable growth over the past few decades, emerging as one of the fastest-growing sectors in food production (FAO, 2020). In 2020, total global aquaculture production reached a record 122.6 million tonnes, valued at approximately USD 281.5 billion (FAO, 2022). This expansion has been largely driven by the rising demand for fish protein and the recognition of aquaculture’s critical role in enhancing food security and nutrition worldwide (FAO, 2022). Asia dominates global aquaculture, accounting for nearly 91.6% of global production, with China contributing over 58% of the total (Verdegem *et al*., 2023). In contrast, Africa accounts for a smaller share of the global aquaculture output at around 2.0% (Verdegem *et al*., 2023). However, the sector is growing steadily in countries like Egypt, Nigeria, and Zambia, which have emerged as key players in the region (Adeleke *et al*., 2020). As with the development of any livestock sector, disease outbreaks are an unavoidable challenge, often necessitating the use of antibiotics and other antimicrobials for outbreak management and control (Newell *et al*., 2010).

This rapid growth has raised concerns about the sustainability of aquaculture practices, particularly in relation to disease management and the widespread use of antibiotics (FAO, 2020). The intensification of aquaculture systems has led to an increase in antimicrobial use (AMU), a common practice to control bacterial diseases and maintain fish health (Bondad Reantaso *et al*., 2023). However, the overuse and misuse of antimicrobials have exacerbated the global challenge of antimicrobial resistance (AMR), a significant health threat that extends beyond human and terrestrial animal populations to aquatic environments (Nadimpalli *et al*., 2018; Deori, Sonowal and Das, 2024). Aquatic systems can serve as reservoirs for resistant pathogens, which can spread through various pathways, including water systems, food chains, and the environment (Nnadozie and Odume, 2019). It is estimated that the aggregate global human, terrestrial and aquatic food animal antibiotic use in 2030 at 236,757 tons (95% UI 145,525–421,426), of which aquaculture constitutes 5.7% but carries the highest use intensity per kilogram of biomass (164.8 mg kg^−1^) (Schar *et al*., 2020).

In many low- and middle-income countries (LMICs), including those in Africa, the challenge of AMR is compounded by weak regulatory frameworks, limited access to veterinary services, and insufficient awareness among farmers (Caudell *et al*., 2020; Mudenda *et al*., 2023). While Zambia’s aquaculture sector is experiencing rapid growth, with fish production from aquaculture facilities increasing from 36,105 metric tonnes (MT) in 2018 to 72,000 MT in 2022, the intensification of farming operations has led to disease outbreaks (Bwalya *et al*., 2020; Siamujompa *et al*., 2023).

In Zambia, fish farmers have reported frequent disease outbreaks in aquaculture facilities, which pose a significant challenge to fish health and production (Bwalya *et al*., 2020; Siamujompa *et al*., 2023). Despite the occurrence of these outbreaks, previous study reported that farmers do not resort to using antibiotics as a treatment measure (Ndashe *et al*., 2023). This is largely due to limited access to veterinary services, and a lack of knowledge about proper antimicrobial use in fish farming (Ndashe *et al*., 2023). Instead, farmers often rely on alternative disease management practices, which may not always be effective in controlling infections, potentially exacerbating fish losses and further threatening the sustainability of their operations (Ndashe *et al*., 2023).

Fish farmers are key actors in the fight against AMR, as their knowledge and practices surrounding antimicrobial use directly impact the development of resistance (Henriksson *et al*., 2018). Understanding their perceptions and behaviors is essential for designing interventions that promote prudent antimicrobial use. However, little is known about how Zambian fish farmers manage the risks associated with AMU and AMR. Addressing this gap is critical for developing policies and programs that reduce antimicrobial misuse in aquaculture and help mitigate the spread of resistant pathogens.

This study aimed to assess the knowledge, attitudes, and practices (KAP) of fish farmers in Zambia regarding AMU and AMR. By evaluating their awareness and behaviors, this research sought to inform the development of strategies to promote responsible AMU and curb the emergence of resistant pathogens in aquaculture systems. The findings will contribute to the global effort to combat AMR while enhancing the sustainability of fish farming in Zambia.

## 2. Materials and Methods

### 2.1. Study Area

This study was conducted in 25 districts across Zambia’s ten provinces, selected based on their high aquaculture activity and fish farming population density. At least two districts were chosen from each province, focusing on regions that significantly contribute to aquaculture development. The selected districts included Gwembe, Sinazongwe, and Siavonga in the Southern Province; Kasempa, Mwinilunga, Solwezi, and Mushindano in the North-Western Province; Kaoma in the Western Province; Chitambo, Chibombo, Mkushi, and Chisamba in the Central Province; Chipata and Petauke in the Eastern Province; Mpulungu, Kasama, and Mungwi in the Northern Province; Mansa and Samfya in the Luapula Province; Ndola in the Copperbelt Province; Chinsali in the Muchinga Province; and Chilanga, Chongwe, Kafue, and Rufunsa in the Lusaka Province [Figure 1].

**Figure 1:**
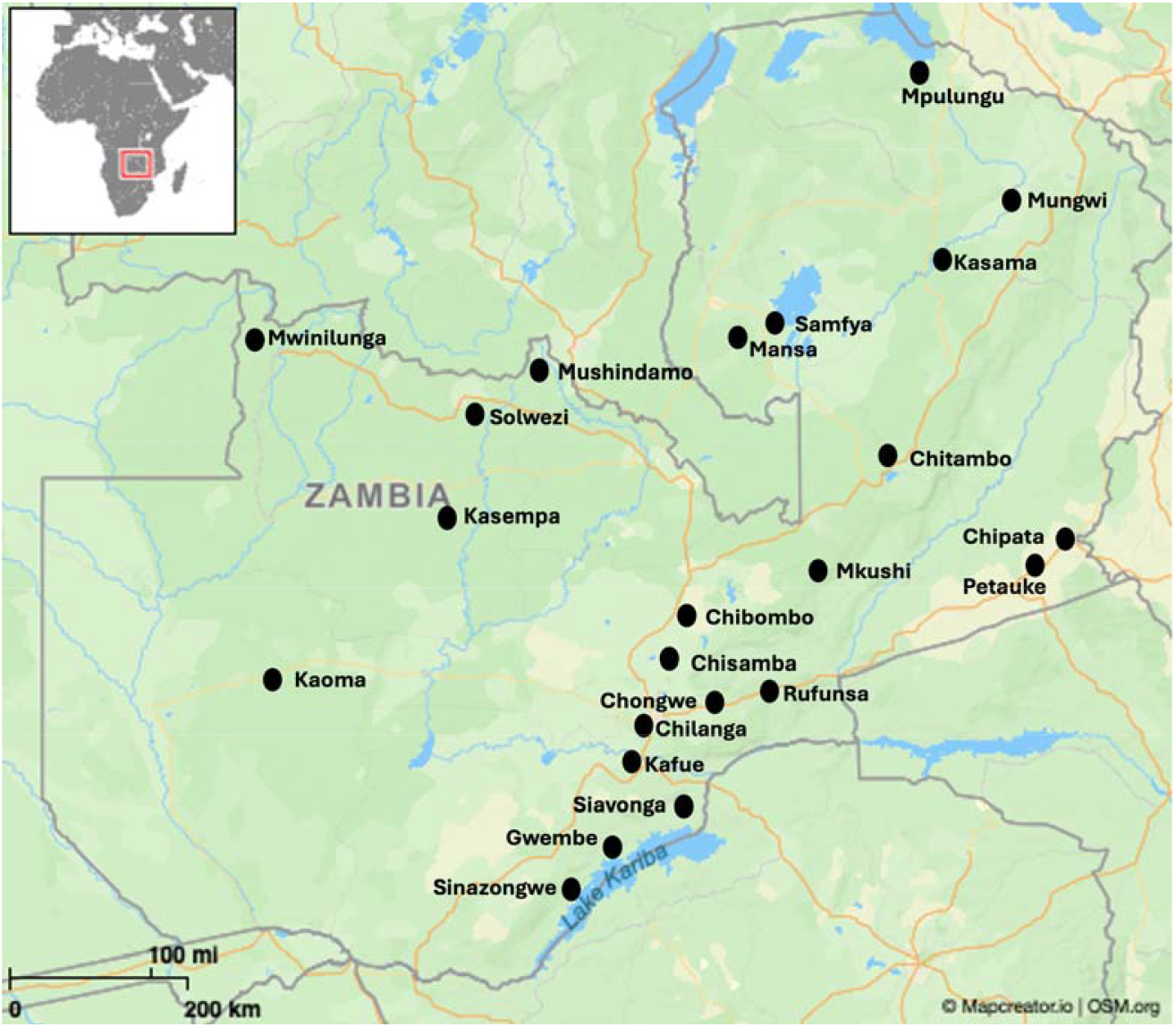
Map showing the districts included in the study.

### 2.2. Study design and Sample Size Determination

A cross-sectional quantitative survey on fish farmers was conducted in different districts of Zambia in September 2024. The sample size was calculated using the Raosoft sample volume calculation method based on a 5% error rate, and 50% response distribution (Raosoft, 2004). As there were no similar types of studies conducted as such in Zambia among fish farmers, we assumed 50% of the respondents would have proper knowledge on AMU, and AMR, added 5% to the sample to address the non-response rate. Thus, a minimum sample size of 170 was necessary for conducting the study.

### 2.3. Data Collection Tool

Data were collected using a structured questionnaire that addressed farmers’ demographics, as well as KAP related to AMU and AMR. Demographic information included farmer age, gender, educational level, work experience, and district location. Questions related to AMU focused on various aspects of aquaculture management, including the type, frequency, source, and storage of antimicrobials, as well as farm management practices concerning school size, fish health, farm biosecurity, clinical signs, and disease occurrence. Knowledge-based questions covered topics such as AMU, antimicrobial withdrawal periods, AMR transmission, and government policies regarding AMU. In the attitudes section, farmers’ perceptions on the safety of AMU and the use of non-prescribed antimicrobials were explored. Finally, practice-related questions sought information on the completion of full courses of AMU, frequency of AMU, and verification of antimicrobial expiration dates. A pre-test of the questionnaire was administered to five farmers, followed by further validation by three experts in AMU and AMR.

### 2.4. Data Collection

The data collection process commenced with virtual meetings with fisheries officers from the selected districts to introduce and familiarize them with the study objectives and the questionnaire. Subsequently, the data collection tool was disseminated to the fisheries officers via WhatsApp. The questionnaire was then distributed to fish farmers through pre-established WhatsApp groups, which were utilized by district fisheries officers for communication with farmers. The questionnaire consisted of two formats: multiple-choice and single-word responses. The survey was administered in English, and responses were electronically recorded using a Google Form (https://forms.gle/imUmkfkMynxnMrtq8), which was accessible for seven days in September 2024. Participation in the study was voluntary, and only farmers who provided informed consent were included in the final dataset.

### 2.5. Statistical Analysis

In this study, both descriptive and inferential statistical analyses were employed. Following the attainment of the required sample size, data were processed using Microsoft Excel 2013 and subsequently analyzed using Datatab software (Team, 2022). The dataset comprised 43 categorical variables, organized into four sections: 10 demographic and socioeconomic variables, 12 knowledge-related variables, 7 attitude-related variables, and 14 practice-related variables.

To assess participants KAP regarding AMU and AMR, a Likert scale was utilized to generate binary dependent variables. Participants who answered at least 7 out of 12 knowledge-based questions correctly were classified as “knowledgeable,” while those answering fewer than 5 were categorized as “not knowledgeable.” Attitudes supporting AMR mitigation were classified as “positive,” while those promoting AMR-contributing behaviors were labeled as “negative.” Practices aimed at reducing AMR were considered “good practice,” while those that increased AMR risk were categorized as “poor practice.”

Binary logistic regression was applied to identify significant predictors of KAP. The analysis was conducted in three stages: first, individual associations between predictor variables and KAP outcomes were assessed; second, descriptive crosstab analyses were used to examine relationships; and third, the impact of demographic and socioeconomic variables on KAP outcomes was expressed through odds ratios.

## 3. Results

### 3.1. Demographic information

The surveyed fish farmers displayed a wide range of demographic and operational characteristics. The largest age groups were 30–39 years (27.72%) and 40–49 years (26.63%), with a male majority (73.37%) [Table 1]. A significant proportion of respondents (65.57%) had received tertiary education, and 84.15% were farm owners. Experience levels varied, with 66.3% having 1–5 years of farming experience, while 22.83% had less than one year of experience [Table 1].

**Table 1:** Demographic information of farmers in study area.

| Variable | n (%) |
| --- | --- |
| Age |  |
| 18 – 29 years | 22 (11.96%) |
| 30 – 39 years | 51 (27.72%) |
| 40 – 49 years | 49 (26.63%) |
| 50 – 59 years | 40 (21.74%) |
| 60 years and above | 22 (11.96%) |
| Gender |  |
| Male | 135 (73.37%) |
| Female | 49 (26.63%) |
| Educational Level |  |
| Primary | 14 (7.65%) |
| Secondary | 44 (24.04%) |
| Tertiary | 120 (65.57%) |
| No formal education | 5 (2.73%) |
| Role on Farm |  |
| Owner | 154 (84.15%) |
| Manager | 25 (13.66%) |
| Worker | 4 (2.19%) |
| Years of involvement in fish farming |  |
| <1 year | 42 (22.83%) |
| 1-5 years | 122 (66.3%) |
| 6-10 years | 12 (6.52%) |
| 11-15 years | 5 (2.72%) |
| ≥16 years | 3 (1.63%) |

| Fish reared by farmers |  |
| --- | --- |
| Three spot tilapia | 86 (43.22%) |
| Nile tilapia | 60 (30.15%) |
| Red breast tilapia | 28 (14.07%) |
| Green head tilapia | 15 (7.54%) |
| Catfish | 10 (5.03%) |
| Production System |  |
| Ponds | 164 (79.61%) |
| Cages | 27 (13.11%) |
| Tanks | 10 (4.85%) |
| Recirculated aquaculture system | 5 (2.43%) |

The most commonly reared species were three-spot tilapia (43.22%) and Nile tilapia (30.15%). Ponds were the predominant production system (79.61%), followed by cages (13.11%), tanks (4.85%), and recirculated aquaculture systems (2.43%) [Table 1].

### 3.2. Socioeconomic information

The primary motivation for fish farming among the surveyed farmers was to generate supplementary income (55.32%), followed by access to free water resources (14.04%) and the perceived ease of farm management (13.62%). Smaller proportions of respondents cited the availability of trained personnel (11.06%) and lower capital requirements (5.96%) as key motivations [Table 2].

**Table 2:** Socioeconomic information of farmers in study area.

| Variable | n(%) |
| --- | --- |
| Reason for starting fish farm |  |
| Supplementary income | 130 (55.32%) |
| Free water resources | 33 (14.04%) |
| Easy to manage | 32 (13.62%) |
| Trained personnel | 26 (11.06%) |
| Less starting capital | 14 (5.96%) |
| Quantity of fish produced per year |  |
| <100 | 11 (5.98%) |
| 100-200 | 15 (8.15%) |
| 201-300 | 13 (7.07%) |
| 301-400 | 5 (2.72%) |
| 401-500 | 14 (7.61%) |
| 501-1000 | 34 (18.48%) |
| 1001-5000 | 42 (22.83%) |
| 5001-10000 | 8 (4.35%) |
| 10001-50000 | 4 (2.17%) |
| 50001-100000 | 2 (1.09%) |
| 100001-150000 | 1 (0.54%) |
| Yet to harvest | 35 (19.02%) |
| Contribution of fish farming to household income |  |
| 0 to 20% | 65 (35.52%) |
| 21 to 40% | 45 (24.59%) |
| 41 to 60% | 24 (13.11%) |
| 61 to 80% | 12 (6.56%) |
| 81 to 100% | 7 (3.83%) |
| I don't know | 3 (1.64%) |
| I have not harvested yet | 27 14.75%) |

Regarding production, 22.83% of farmers reported annual yields ranging from 1,001–5,000 kg, while 19.02% had not yet harvested fish, suggesting they were in the early stages of farming. Smaller production ranges included fewer than 100 kg (5.98%) and 501–1,000 kg (18.48%). A small proportion of farmers (2.17%) reported production levels above 10,000 kg [Table 2].

Fish farming contributed modestly to household incomes, with 35.52% of farmers reporting a contribution of 0–20%, and 24.59% indicating a contribution of 21–40%. Only 6.56% reported contributions of 61–80%, and 3.83% indicated contributions of 81–100%. Additionally, 14.75% of farmers had not yet harvested fish, limiting their ability to assess the financial impact of their operations [Table 2].

### 3.3. Knowledge Assessment

The knowledge assessment of fish farmers regarding AMU and AMR in aquaculture revealed varying levels of awareness. A majority of farmers (71.89%) were aware of antibiotics, with amoxicillin being the most recognized example (55.68%). However, 27.03% of respondents were uncertain. Veterinarians were identified by 31.89% as the primary authority for prescribing antibiotics, followed by fisheries officers (27.03%), though 32.97% were unsure.

Awareness of the risks associated with antibiotics was moderate. Specifically, 47.57% of farmers recognized the potential side effects of antibiotics on fish and the environment, and 55.23% were aware that antibiotics could enter the human body through fish consumption. Additionally, 52.69% understood the concept of antibiotic residues, and 65.05% had heard of antibiotic resistance, with 54.39% acknowledging the potential for pathogen transmission to humans through fish products.

Regarding antibiotic resistance, 63.44% of farmers believed it could be mitigated by avoiding antibiotic overuse, while 81.18% recognized the harmful effects of antimicrobial misuse on animal, human, and environmental health. Biosecurity and hygiene were widely regarded as critical, with 76.34% acknowledging the importance of biosecurity measures and 67.74% emphasizing the role of proper hygiene practices in reducing antibiotic resistance.

**Table 1:**
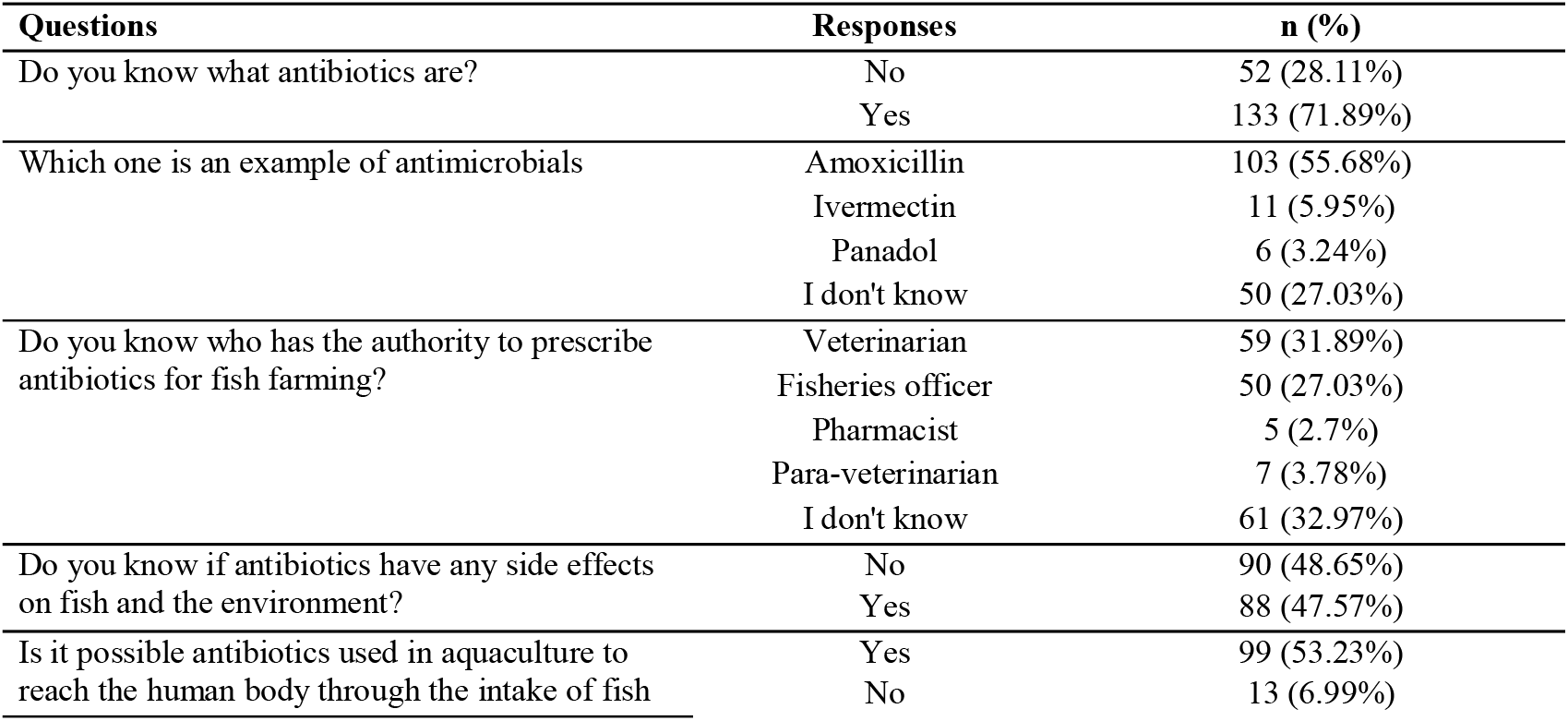
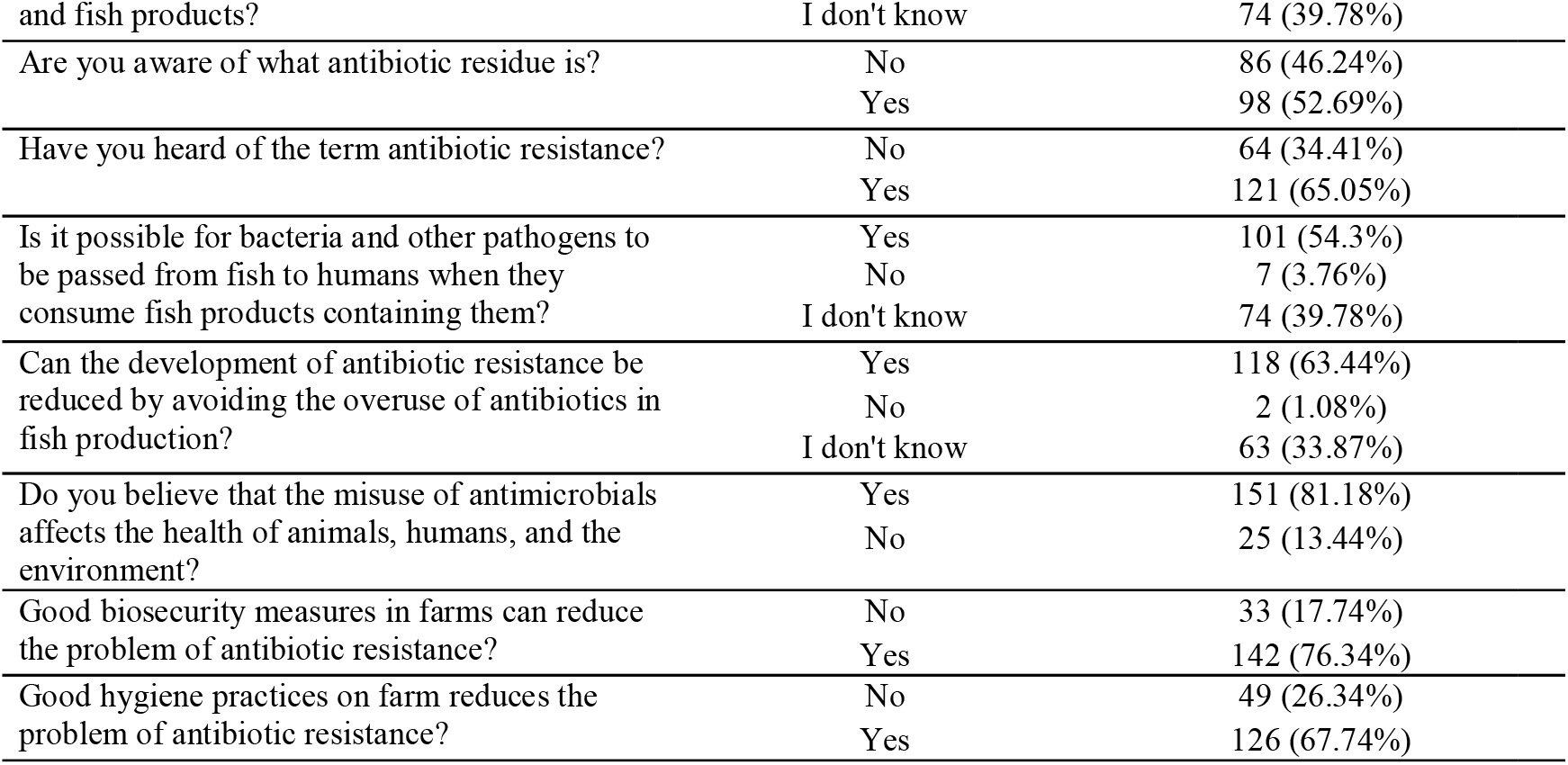
Knowledge assessment of farmers in study area.

A significant majority of farmers (82.6%) exhibited insufficient knowledge regarding proper antimicrobial use (AMU) and antimicrobial resistance (AMR), with only 17.4% demonstrating adequate understanding [Figure 2]. This finding highlights a considerable knowledge gap, emphasizing the need for targeted educational and awareness initiatives to mitigate AMR risks in aquaculture.

**Figure 2.**
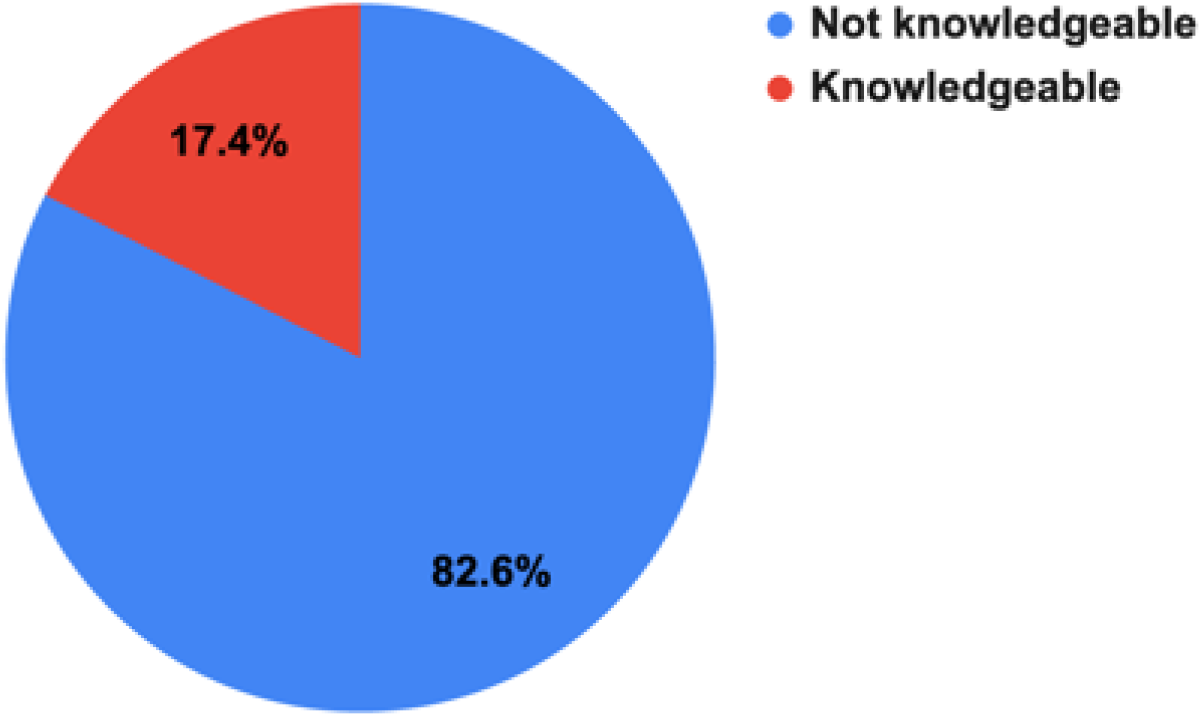
graph showing the percent of farmers Knowledgeable on not in AMU and AMR.

### 3.4. Attitude Assessment

The majority of farmers (76.34%) believe that fish should be treated when clinical signs or mortality occur, while 10.75% disagree and 11.29% are uncertain. Most farmers (83.87%) agree on the importance of seeking veterinary advice prior to using antibiotics, with only 1.61% disagreeing and 13.98% unsure. Additionally, 75.27% support the regulation of antibiotic use by authorities, although 22.58% are uncertain and 1.08% disagree.

A strong majority (89.25%) believe that education and public awareness are crucial for reducing antibiotic resistance, and 63.44% recognize the environmental risks associated with antibiotic use, though 28.49% remain uncertain. Furthermore, 78.49% consider antibiotic resistance to be a public health concern, and 81.18% acknowledge their role in mitigating it. These findings indicate a generally positive attitude toward responsible antibiotic use, regulation, and awareness initiatives [Table 4].

**Table 3:** A Logistic Regression Analysis of Influence of Demographic and Farm Characteristics on Farmers’ Knowledge, Attitudes, and Practices Regarding Antimicrobial Use and Resistance.

| Variables |  | Knowledge |  |  | Attitude |  |  | Practice |  |  |
| --- | --- | --- | --- | --- | --- | --- | --- | --- | --- | --- |
|  |  | OR | 95% CI, | p | OR | 95% CI, | p | OR | 95% CI, | p |
| Age of the farmers (Years) | 18 – 29 |  | Ref |  |  | Ref |  |  | Ref |  |
|  | 30 – 39 | 5.78 | 0.7 - 47.83 | 0.104 | **2.88** | **1.03 - 8.09** | **0.045** | 1.66 | 0.59 - 4.71 | 0.339 |
|  | 40 – 49 | 4.1 | 0.48 - 34.98 | 0.197 | 2.72 | 0.97 - 7.67 | 0.058 | 2.92 | 0.96 - 8.94 | 0.06 |
|  | 50 – 59 | 4.45 | 0.51 - 38.84 | 0.176 | 1.4 | 0.49 - 4 | 0.53 | 3.92 | **1.16 - 13.22** | **0.027** |
|  | >60 | 6.56 | 0.7 - 61.86 | 0.1 | 1 | 0.31 - 3.28 | 1 | 4.15 | 0.94 - 18.41 | 0.061 |
| Gender | Female |  | Ref |  |  | Ref |  |  | Ref |  |
|  | Male | 1.86 | 0.83 - 4.17 | 0.13 | 0.81 | 0.41 - 1.58 | 0.534 | 0.88 | 0.4 - 1.92 | 0.751 |
| Educational Level | No formal education |  | Ref |  |  | Ref |  |  | Ref |  |
|  | Primary | 1 | 0.07 - 13.37 | 1 | 4.89 | 0.68 - 34.97 | 0.114 | 4.5 | 0.54 - 37.38 | 0.164 |
|  | Secondary | 0.77 | 0.08 - 7.77 | 0.824 | 1.6 | 0.32 - 8.01 | 0.567 | 1.55 | 0.31 - 7.91 | 0.596 |
|  | Tertiary | 1.52 | 0.17 - 13.19 | 0.706 | 2.12 | 0.45 - 9.89 | 0.341 | 2.94 | 0.62 - 14.02 | 0.177 |
| Role on Farm | Worker |  | Ref |  |  | Ref |  |  | Ref |  |
|  | Owner | **0.19** | **0.04 - 0.98** | **0.047** | 0 | 0 - ∞ | 0.998 | 1.74 | 0.3 - 9.88 | 0.535 |
|  | Manager | 0.25 | 0.04 - 1.63 | 0.148 | 0 | 0 - ∞ | 0.998 | 1.21 | 0.18 - 8.22 | 0.842 |
| Experience in farming | <1year |  | Ref |  |  | Ref |  |  | Ref |  |
|  | 1-5 years | 1.14 | 0.45 - 2.9 | 0.779 | 2.46 | **1.21 - 5.01** | **0.013** | 2.62 | **1.19 - 5.76** | **0.017** |
|  | 6-10 years | 1.03 | 0.18 - 5.75 | 0.974 | 3.10 *10<sup>9</sup> | 0 - ∞ | 0.998 | 1.67 | 0.39 - 7.11 | 0.49 |
|  | 11-15 years | 1.29 | 0.12 - 13.3 | 0.833 | 2.08 | 0.32 - 13.78 | 0.446 | 0.37 | 0.06 - 2.47 | 0.305 |
|  | ≥16 years | 0 | 0 - ∞ | 0.998 | 0 | 0 - ∞ | 0.999 | 1.11 | 0.09 - 13.3 | 0.934 |
| Annual Production Capacity (Kg) | <100 |  | Ref |  |  | Ref |  |  | Ref |  |
|  | 101-200 | 1 | 0 - ∞ | 1 | 0.38 | 0.08 - 1.9 | 0.239 | 4 | 0.37 - 43.14 | 0.253 |
|  | 201-300 | 1 | 0 - ∞ | 1 | 1.9 | 0.32 - 11.31 | 0.478 | 108 | **5.92 - 1969.64** | **0.002** |
|  | 301-400 | 1 | 0 - ∞ | 1 | 2.29 | 0.19 - 28.19 | 0.519 | 6 | 0.39 - 92.28 | 0.199 |
|  | 401-500 | 3.55*10<sup>9</sup> | 0 - ∞ | 0.999 | 0.57 | 0.11 - 2.87 | 0.497 | 16.2 | 1.56 - 167.75 | 0.02 |
|  | 501-1000 | 1.30 * 10<sup>10</sup> | 0 - ∞ | 0.999 | 3.31 | 0.7 - 15.65 | 0.13 | 93 | **8.59 - 1006.65** | **<.001** |
|  | 1001-5000 | 5.58 * 10<sup>9</sup> | 0 - ∞ | 0.999 | 1.27 | 0.32 - 5.13 | 0.732 | 66.6 | **6.9 - 642.9** | **<.001** |
|  | 5001-10000 | 1 | 0 - ∞ | 1 | 1.71 | 0.23 - 12.89 | 0.601 | 9 | 0.75 - 108.32 | 0.083 |
|  | 10001-50000 | 1 | 0 - ∞ | 1 | 0.18 | **0.04 - 0.78** | **0.021** | 102 | **9.44 - 1101.54** | **<.001** |
|  | 50001-100000 | 1 | 0 - ∞ | 1 | 1.71 | 0.13 - 22.51 | 0.682 | 1.19 *10<sup>10</sup> | 0 - ∞ | 0.998 |
|  | 100001-150000 | 1 | 0 - ∞ | 1 | 1.73 * 10<sup>8</sup> | 0 - ∞ | 0.999 | 1.19 *10<sup>10</sup> | 0 - ∞ | 0.999 |
|  | Yet to harvest | 1 | 0 - ∞ | 1 | 1.73 * 10<sup>8</sup> | 0 - ∞ | 0.999 | 0 | 0 - ∞ | 0.999 |

**Table 4:** Attitude assessment of farmers towards issues of AMU and AMR in the study area.

| Questions | Responses | n (%) |
| --- | --- | --- |
| Do you think it is necessary to treat the fish once you have noticed clinical signs or mortality? | Agree | 142 (76.34%) |
|  | Disagree | 20 (10.75%) |
|  | I don't know | 21 (11.29%) |
| Do you think seeking advice from a veterinarian is necessary before using antibiotics? | Agree | 156 (83.87%) |
|  | Disagree | 3 (1.61%) |
|  | I don't know | 26 (13.98%) |
| Do you believe that regulating antibiotic use by the Zambia Medicines Regulatory Authority (ZAMRA) or Veterinary Council of Zambia (VCZ) can reduce the irrational use of antibiotics in fish farming? | Agree | 140 (75.27%) |
|  | Disagree | 2 (1.08%) |
|  | I don't know | 42 (22.58%) |
| Do you think education and public awareness can help reduce the problem of antibiotic resistance? | Agree | 166 (89.25%) |
|  | Disagree | 1 (0.54%) |
|  | I don't know | 19 (10.22%) |
| Do you believe that the use of antibiotics in fish farming can affect environmental health? | Agree | 118 (63.44%) |
|  | Disagree | 14 (7.53%) |
|  | I don't know | 53 (28.49%) |
| Do you think antibiotic resistance is a public health concern that should be prevented? | Agree | 146 (78.49%) |
|  | Disagree | 2 (1.08%) |
|  | I don't know | 37 (19.89%) |
| Do you think farmers have a critical role to play to reduce antibiotic resistance? | Agree | 151 (81.18%) |
|  | Disagree | 1 (0.54%) |
|  | I don't know | 33 (17.74%) |

The assessment of farmers’ attitudes revealed that 60.5% exhibited a positive attitude toward responsible antibiotic use (AMU) and antimicrobial resistance (AMR), while 39.5% demonstrated a negative attitude. This suggests that a significant proportion of the farming community remains unaware, indifferent, or resistant to proper AMU and AMR practices, highlighting the need for ongoing efforts to enhance awareness and improve attitudes within the sector [Figure 3].

**Figure 3.**
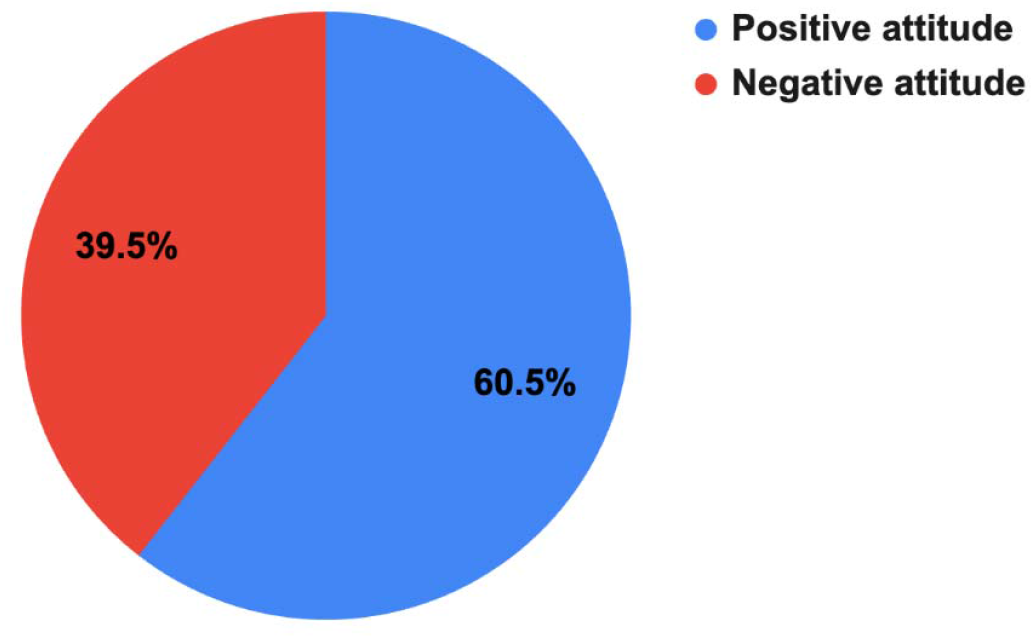
graph showing the percent of farmers with positive and negative attitude in AMU and AMR.

### 3.5 Practice Assessment

The majority of farmers (94.09%) reported not using antibiotics on their fish farms, while 3.23% confirmed antibiotic use, primarily for treatment (2.69%) and less commonly for both treatment and growth promotion (1.62%). Specific antibiotics cited included coarse salt (2.15%), formalin (1.08%), penicillin (0.54%), and amoxicillin (0.54%). Most farmers (84.29%) had never purchased antibiotics, with those who did primarily sourcing them from agrovets (6.44%).

Only 9.89% obtained prescriptions from veterinarians, while 43.01% administered antibiotics without adhering to guidelines. Additionally, 50.27% of farmers did not submit fish samples for veterinary testing, and 74.19% reported no antibiotic use. Proper disposal of antibiotic containers was practiced by 6.99% of farmers, while others buried (2.15%) or discarded them in open pits (1.61%). For expired antibiotics, 1.61% burned them, 4.84% discarded them in open pits, and 0.54% shared them with others.

When fish exhibited symptoms, 42.47% consulted veterinarians, 12.37% isolated and treated sick fish, and 9.68% treated all fish. These findings indicate limited antibiotic use among farmers, but also reveal significant gaps in proper administration, disposal practices, and veterinary engagement [Table 5].

**Table 5:** Assessment of fish farmers in study area in practices towards AMU and AMR.

| Questions | Responses | n (%) |
| --- | --- | --- |
| Do you use antibiotics on your fish farm? | No | 175 (94.09%) |
|  | Yes | 6 (3.23%) |
| For what purpose do you use antibiotics most? | I don't use antibiotics | 107 (57.52%) |
|  | Treatment only | 5 (2.69%) |
|  | Treatment and grow promoter | 3 (1.62%) |
|  | Growth promoter | 1 (0.54%) |
| Give an example of antibiotics you have used on your fish farm | Coarse Salt | 4 (2.16%) |
|  | Formalin | 2 (1.08%) |
|  | Penicillin | 1 (0.54%) |
|  | Amoxicillin | 1 (0.54%) |
|  | I don't use antibiotics | 73 (39.44%) |
| Where do you usually get your antibiotics from? | Agrovet | 6 (4.29%) |
|  | Chemist/pharmacy | 4 (2.86%) |
|  | Private Veterinary Clinic | 5 (3.57%) |
|  | Government Veterinary Clinic | 4 (2.86%) |
|  | Feed distributor | 3 (2.14%) |
|  | I have never purchased | 118 (84.29%) |
| Are the antibiotics you used as effective as before? | Yes | 6 (3.23%) |
|  | No | 4 (2.15%) |
|  | I don't use antibiotics | 129 (69.35%) |
|  | I don't know | 4 (2.15%) |
| Who provides you with the prescriptions for the antibiotics? | Veterinarian | 18 (9.89%) |
|  | Para-veterinarian | 4 (2.78%) |
|  | Self-treatment follow instructions | 6 (4.17%) |
|  | Other farmers/friends | 5 (3.13%) |
|  | I don't use antibiotics | 60 (34.48%) |
| Do you refer to guidelines or calculate the dose while administering the antibiotics? | No | 80 (43.01%) |
|  | Yes | 28 (15.05%) |
| Do you wait the required time after using antibiotics before eating or selling fish? | I don't use antibiotics | 132 (70.97%) |
|  | Yes | 11 (5.91%) |
|  | No | 3 (1.61%) |
| Do you submit samples of diseased fish to a veterinary laboratory? | No | 93 (50.27%) |
|  | Yes | 28 (15.14%) |
| How long do you continue using antibiotics once you start? | 1-2 days | 1 (0.54%) |
|  | 3-4 days | 1 (0.54%) |
|  | 5-7 days | 3 (1.61%) |
|  | Stop medication when symptoms subside | 5 (2.69%) |
|  | I don't use antibiotics | 138 (74.19%) |
| Do you keep a record of antibiotics you used earlier on the farm? | Yes | 7 (3.76%) |
|  | No | 3 (1.61%) |
|  | I don't know | 1 (0.54%) |
|  | I don't use antibiotics | 138 (74.19%) |
| When one or a few fish display symptoms what do you do? | Call veterinarian/paravet | 79 (42.47%) |
|  | Call another farmer for help | 25 (13.44%) |
|  | Isolate sick ones and treat them | 23 (12.37%) |
|  | Treat all | 18 (9.68%) |
| How do you dispose containers after the use of antibiotics | Burn | 13 (6.99%) |
|  | Bury them | 4 (2.15%) |
|  | Throw in the trash to be taken to dumpsite | 3 (1.61%) |
|  | Throw them in a pit | 3 (1.61%) |
|  | I don't treat my fish | 125 (67.20%) |
| What do you usually do with expired antibiotics? | Give to friend/neighbour/fellow farmer to use | 1 (0.54%) |
|  | Bury them | 3 (1.61%) |
|  | Burn | 11 (5.91%) |
|  | Throw them in an open pit | 9 (4.84%) |
|  | I don't treat my fish | 126 (67.74%) |

The assessment of practice levels revealed that 76.5% of farmers adhere to good practices, including responsible antibiotic use, consultation with veterinary professionals, and efforts to mitigate antimicrobial resistance (AMR) risks. However, 23.5% engage in poor practices, indicating non-compliance with established protocols, which may contribute to the development of AMR. These findings underscore the need for continued education and targeted interventions to promote the widespread adoption of responsible antibiotic use practices [Figure 4].

**Figure 4.**
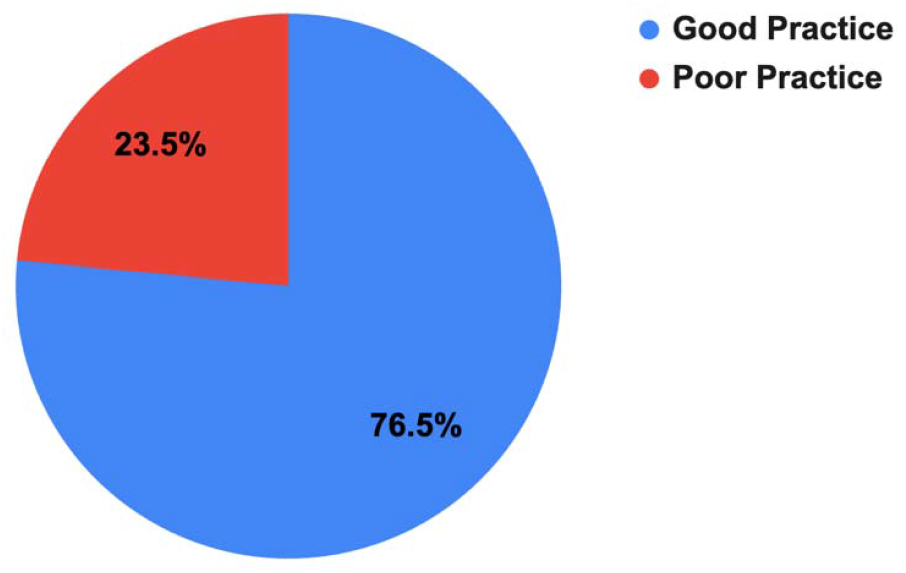
graph showing the percent of farmers with good and poor practices in AMU and AMR.

### 3.6. Clinical Signs of Diseased fish observed by farmers

The KAP survey identified skin wounds as the most commonly reported clinical sign of fish diseases (47.85%), followed by unusual swimming behavior (43.55%), loss of appetite (37.63%), and gasping for air at the surface (37.10%). Other reported signs included white or cloudy skin spots (26.34%), reddish skin spots (22.04%), whitish gills (18.28%), sunken or protruding eyes (17.74%), and whitish eyes (13.98%). Excessive mucus or slime on the body was the least reported clinical sign (12.37%) [Figure 5].

**Figure 5:**
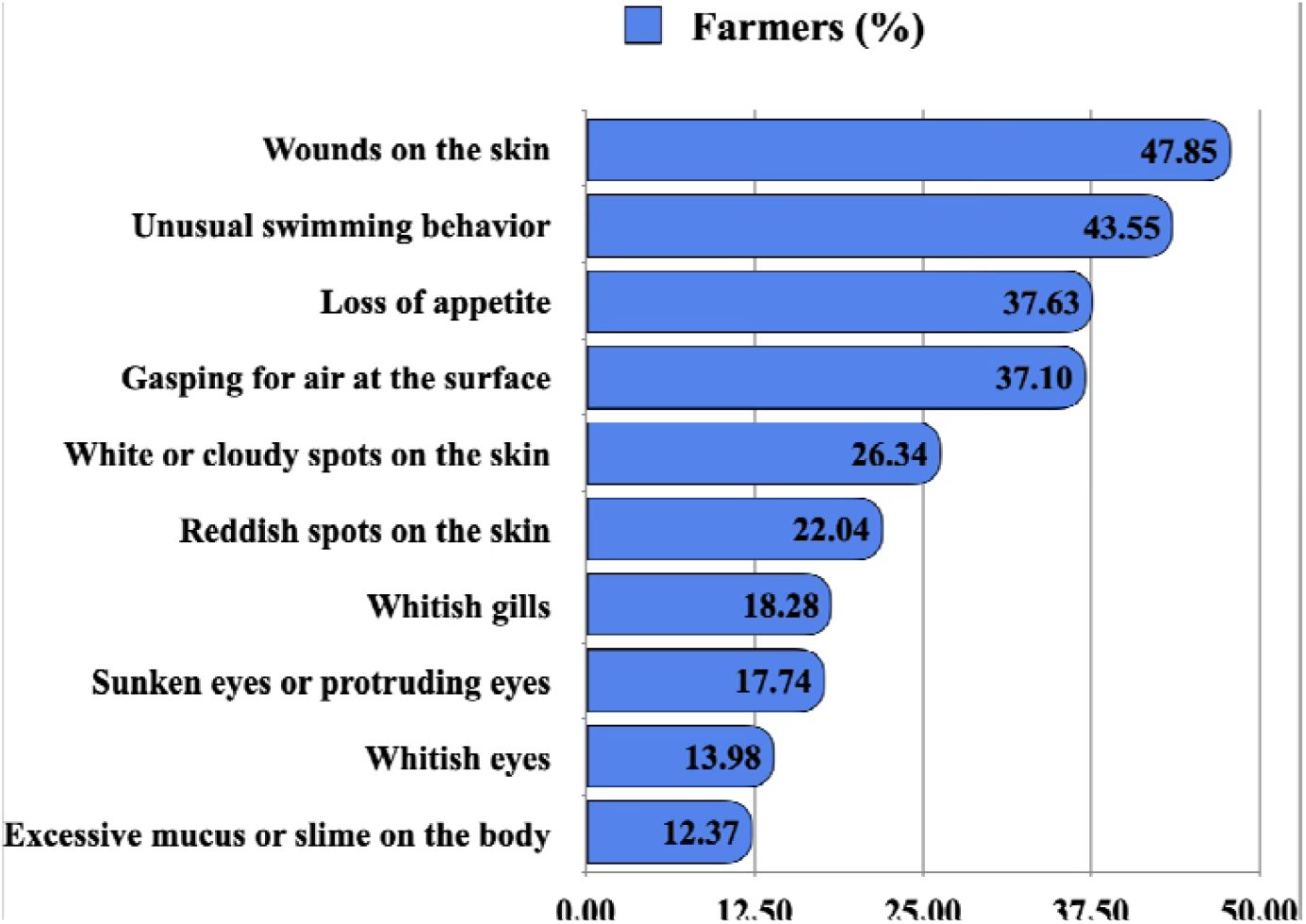
Clinical signs of fish reported by farmers in study area.

### 3.7. Influence of Demographic and Operational Factors on Farmers’ Knowledge, Attitudes, and Practices Regarding AMU and AMR

Statistical analysis of the KAP regarding AMU and AMR revealed key demographic and operational trends. While age, gender, education, and farm roles showed no significant differences in knowledge, farmers aged over 60 (23.81%) and those with tertiary education (20.17%) exhibited higher knowledge levels. Knowledge was significantly associated with annual production capacity, with farmers producing 501–1000 kg demonstrating better knowledge (p < 0.001).

Positive attitudes were more prevalent among farmers aged 30–39 (70.59%) and those with 6–10 years of experience (p = 0.002). Larger producers also exhibited significantly more positive attitudes (p < 0.001). Regarding practices, farmers with larger production capacities (501–1000 kg) and 1–5 years of experience displayed better practices (p-values < 0.001 and 0.039, respectively). While age showed borderline significance (p = 0.097), farmers with higher production capacities were more likely to follow good practices (91.18%).

Overall, larger producers and more experienced farmers demonstrated better KAP concerning AMU and AMR, underscoring the need for targeted educational interventions aimed at smaller-scale and less experienced farmers.

**Table 2:** Test of the statistical significances of the variations in the respondents’ knowledge of AMU and AMR by their characteristics.

| Variables |  | Knowledge |  | p-Value | Attitude |  | p-Value | Practice |  | p-Value |
| --- | --- | --- | --- | --- | --- | --- | --- | --- | --- | --- |
|  |  | Knowledgeable | Not knowledgeable |  | Positive attitude | Negative attitude |  | Good Practice | Poor Practice |  |
| Age of the farmers (Years) | 18 – 29 | 1 (4.55%) | 21 (95.45%) | 0.438 | 10 (45.45%) | 12 (54.55%) | 0.069 | 13 (59.09%) | 9 (40.91%) | 0.097 |
|  | 30 – 39 | 11 (21.57%) | 40 (78.43%) |  | 36 (70.59%) | 15 (29.41%) |  | 36 (70.59%) | 15 (29.41%) |  |
|  | 40 – 49 | 8 (16.33%) | 41 (83.67%) |  | 34 (69.39%) | 15 (30.61%) |  | 38 (80.85%) | 9 (19.15%) |  |
|  | 50 – 59 | 7 (17.50%) | 33 (82.50%) |  | 21 (53.85%) | 18 (46.15%) |  | 34 (85.00%) | 6 (15.00%) |  |
|  | >60 | 5 (23.81%) | 16 (76.19%) |  | 10 (45.45%) | 12 (54.55%) |  | 18 (85.71%) | 3 (14.29%) |  |
| Gender | Female | 12 (25.00%) | 36 (75.00%) | 0.111 | 31 (63.27%) | 18 (36.73%) | 0.598 | 39 (79.59%) | 10 (20.41%) | 0.587 |
|  | Male | 20 (14.82%) | 115 (85.19%) |  | 79 (58.96%) | 55 (41.04%) |  | 100 (75.76%) | 32 (24.24%) |  |
| Educational Level | No formal education | 1 (20.00%) | 4 (80.00%) | 0.604 | 2 (40.00%) | 3 (60.00%) | 0.324 | 3 (60.00%) | 2 (40.00%) | 0.256 |
|  | Primary | 2 (14.29%) | 12 (85.71%) |  | 11 (78.57%) | 3 (21.43%) |  | 12 (85.71%) | 2 (14.29%) |  |
|  | Secondary | 5 (11.36%) | 39 (88.64%) |  | 24 (54.54%) | 20 (45.46%) |  | 29 (67.44%) | 14 (32.56%) |  |
|  | Tertiary | 24 (20.17%) | 95 (79.83%) |  | 73 (61.34%) | 46 (38.66%) |  | 94 (79.66%) | 24 (20.34%) |  |
| Role on Farm | Manager | 5 (20.00%) | 20 (80.00%) | 0.18 | 15 (62.50%) | 9 (37.50%) | 0.237 | 17 (70.83%) | 7 (29.17%) | 0.763 |
|  | Owner | 24 (15.69%) | 129 (84.31%) |  | 90 (58.44%) | 64 (41.56%) |  | 118 (77.63%) | 34 (22.37%) |  |
|  | Worker | 2 (50.00%) | 2 (50.00%) |  | 4 (100.00%) | 0 (0.00%) |  | 3 (75.00%) | 1 (25.00%) |  |
| Experience in farming | <1year | 7 (16.67%) | 35 (83.33%) | 0.948 | 18 (42.86%) | 24 (57.14%) | 0.002 | 26 (63.41%) | 15 (36.59%) | 0.039 |
|  | 1-5 years | 22 (18.18%) | 99 (81.82%) |  | 78 (63.93%) | 44 (36.07%) |  | 99 (82.50%) | 21 (17.50%) |  |
|  | 6-10 years | 2 (16.67%) | 10 (83.33%) |  | 12 (100.00%) | 0 (0.00%) |  | 9 (75.00%) | 3 (25.00%) |  |
|  | 11-15 years | 1 (20.00%) | 4 (80.00%) |  | 3 (60.00%) | 2 (40.00%) |  | 2 (40.00%) | 3 (60.00%) |  |
|  | ≥16 years | 0 (0.00%) | 3 (100.00%) |  | 0 (0.00%) | 2 (100.00%) |  | 2 (66.67%) | 1 (33.33%) |  |
| Annual Production Capacity (Kg) | <100 | 0 (0.00%) | 11 (100.00%) | <.001 | 7 (63.64%) | 4 (36.36%) | <.001 | 1 (10.00%) | 9 (90.00%) | <.001 |
|  | 101-200 | 0 (0.00%) | 15 (100.00%) |  | 6 (40.00%) | 9 (60.00%) |  | 4 (30.77%) | 9 (69.23%) |  |
|  | 201-300 | 0 (0.00%) | 13 (100.00%) |  | 10 (76.92%) | 3 (23.08%) |  | 12 (92.31%) | 1 (7.69%) |  |
|  | 301-400 | 0 (0.00%) | 5 (100.00%) |  | 4 (80.00%) | 1 (20.00%) |  | 2 (40.00%) | 3 (60.00%) |  |
|  | 401-500 | 3 (21.43%) | 11 (78.57%) |  | 7 (50.00%) | 7 (50.00%) |  | 9 (64.29%) | 5 (35.71%) |  |
|  | 501-1000 | 17 (50.00%) | 17 (50.00%) |  | 29 (85.29%) | 5 (14.71%) |  | 31 (91.18%) | 3 (8.82%) |  |
|  | 1001-5000 | 12 (30.00%) | 28 (70.00%) |  | 29 (69.05%) | 13 (30.95%) |  | 37 (88.09%) | 5 (11.91%) |  |
|  | 5001-10000 | 0 (0.00%) | 8 (100.00%) |  | 6 (75.00%) | 2 (25.00%) |  | 4 (50.00%) | 4 (50.00%) |  |
|  | 10001-50000 | 0 (0.00%) | 4 (100.00%) |  | 3 (75.00%) | 1 (25.00%) |  | 4 (100.00%) | 0 (0.00%) |  |
|  | 50001-100000 | 0 (0.00%) | 2 (100.00%) |  | 1 (100.00%) | 0 (0.00%) |  | 2 (100.00%) | 0 (0.00%) |  |
|  | 100001-150000 | 0 (0.00%) | 1 (100.00%) |  | 1 (100.00%) | 0 (0.00%) |  | 0 (0.00%) | 1 (100.00%) |  |
|  | Yet to harvest | 0 (0.00%) | 37 (100.00%) |  | 9 (24.32%) | 28 (75.68%) |  | 34 (91.89%) | 3 (8.11%) |  |

### 3.8. Determinants of Farmers’ Knowledge, Attitudes, and Practices Toward AMU and AMR: Insights from Logistic Regression Analysis

Logistic regression analysis identified key factors influencing farmers’ KAP regarding AMU and AMR. Age was a significant determinant, with farmers aged 30–39 being more likely to exhibit positive attitudes (OR = 2.88, p = 0.045), while those aged 40–49 demonstrated better practices (OR = 3.92, p = 0.027). Experience also played a crucial role, as farmers with 1–5 years of experience displayed significantly better attitudes (OR = 2.46, p = 0.013) and practices (OR = 2.62, p = 0.017) compared to less experienced farmers.

Annual production capacity strongly influenced KAP outcomes, with farmers producing 201– 500 kg being much more likely to adopt good practices (OR = 108.0, p = 0.002; OR = 93.0, p< 0.001), indicating better access to resources and training. Interestingly, farm workers were more knowledgeable than farm owners (OR = 0.19, p = 0.047), likely due to their direct involvement in daily farm operations. However, gender and education level were not significantly associated with KAP.

## 4. Discussion

The study provides crucial insights into the demographic and operational characteristics of fish farmers in Zambia, alongside their KAP regarding AMU and AMR. These findings highlight the significant gaps in awareness and responsible practices, particularly concerning the use of antibiotics in aquaculture. When compared to similar studies conducted globally, it is evident that the challenges Zambia faces in addressing AMU and AMR are not unique, but reflective of trends observed across aquaculture systems in many developing countries (Bondad Reantaso *et al*., 2023; Caputo *et al*., 2023).

The demographic profile of fish farmers in this study reveals a diverse range of ages, educational backgrounds, and levels of experience. The majority of respondents were middle-aged, with most having attained tertiary education, suggesting that aquaculture is attracting educated individuals. However, despite this high level of formal education, many farmers still exhibit limited knowledge of AMU and AMR, a trend that has been observed globally (Jia *et al*., 2017; Pham-Duc *et al*., 2019; Chowdhury *et al*., 2022; Dandi *et al*., 2024). In countries such as Bangladesh and Vietnam, where aquaculture plays a significant role in the economy, studies have found that even farmers with formal education often lack specific training on antibiotic use and biosecurity (Pham-Duc *et al*., 2019; Chowdhury *et al*., 2022). This emphasizes that general education alone is insufficient to equip farmers with the necessary skills and knowledge to manage AMU and AMR responsibly.

The lack of significant difference in knowledge based on educational background in this study suggests that while formal education is important, it is specialized training that truly influences AMU and AMR awareness. The finding that farmers with larger production capacities had better knowledge and practices supports this argument, as they likely have more access to technical training, veterinary services, and resources that improve their management practices. This is consistent with findings from Ghanat, where larger commercial fish farms tend to follow better AMU practices due to their capacity to engage professional support and access to advanced aquaculture systems (Dandi *et al*., 2024).

The knowledge assessment in this study paints a concerning picture, with only 17.4% of farmers considered knowledgeable about AMU and AMR. While 71.79% were aware of what antibiotics are, fewer had a clear understanding of antimicrobial resistance, its causes, and its consequences for both fish farming and human health. A comparison with studies from Southeast Asia, such as those by Pham-Duc et al. (2019), shows similar trends where farmers recognize the term “antibiotics” but have limited understanding of their responsible use, and the long-term impacts of overuse on public health and environmental ecosystems (Pham-Duc *et al*., 2019).

The widespread knowledge gap observed in Zambia aligns with the situation in other developing nations, where aquaculture systems are expanding rapidly, but regulatory frameworks and education have not kept pace. For example, in Vietnam, Strom et al. (2019) reported that while many fish farmers were aware of antibiotic use, they lacked understanding of how antibiotic resistance develops and spreads through aquaculture systems (Ström *et al*., 2019). This points to a global need for more targeted education and training programs focused specifically on AMU and AMR in aquaculture, rather than general agricultural practices.

The majority of farmers (60.5%) exhibited a positive attitude toward responsible AMU and AMR practices, indicating an openness to adopting better practices if given the right information and support. This aligns with findings from studies in countries such as Thailand and India, where farmers generally expressed positive attitudes toward seeking veterinary advice and following recommended treatments, but often lacked access to reliable services and faced economic constraints that limited their ability to implement best practices (Baoprasertkul, Somsiri and Boonyawiwat, 2012; Kumaran *et al*., 2012).

A key observation in this study is the significant role of experience in shaping attitudes. Farmers with 1–5 years of experience were more likely to have positive attitudes towards AMU and AMR compared to less experienced counterparts. This suggests that practical exposure to the challenges of fish farming, including disease management, may foster a greater appreciation for the importance of responsible antibiotic use. It also highlights the need for ongoing support for new entrants into the industry, to ensure they develop positive practices early in their farming careers.

The findings on farmers’ practices regarding AMU in Zambia reveal a stark contrast between awareness and action. Despite a strong recognition of the importance of veterinary guidance (with 83.87% agreeing on the need for veterinary consultation), only 15.05% of farmers adhered to dosing guidelines, and most did not submit diseased fish samples to laboratories for diagnosis. This gap between belief and practice is a common issue in aquaculture. Similar studies in India have shown that while farmers understand the need for proper antibiotic use, they often fail to comply with best practices due to logistical, financial, or accessibility issues (Patil *et al*., 2022).

The predominant use of ponds as the main production system (79.61%) and the low adoption of more intensive systems, such as recirculating aquaculture systems (2.43%), may contribute to the challenges of AMU management. Pond-based systems tend to have lower levels of biosecurity, which can lead to higher disease outbreaks and the consequent increased use of antibiotics. This is a trend observed in other regions as well, where intensive systems are associated with better biosecurity and lower dependency on antibiotics (Flores *et al*., 2015; Bera *et al*., 2018).

Biosecurity measures, such as proper disposal of antibiotic containers and isolation of sick fish, were poorly practiced, with only a small proportion of farmers following these protocols. This is concerning given the global emphasis on biosecurity as a key strategy to reduce the spread of AMR in aquaculture. Studies from developed countries, such as Norway and Scotland, where biosecurity protocols are strictly enforced, show much lower levels of antibiotic use due to better disease prevention practices (Lillehaug, Santi and Østvik, 2015; Murray and Gubbins, 2016; Subasinghe *et al*., 2023).

One of the most striking findings in this study is that 94.09% of farmers reported not using antibiotics on their farms, while those who did primarily used them for treatment rather than growth promotion. While this may initially appear positive, it raises questions about the underreporting of antibiotic use or the potential use of unregulated, non-approved products. In countries like Zambia, where regulatory oversight is still developing, there is often a lack of accurate data on antibiotic use, as informal markets and the lack of proper record-keeping make monitoring difficult. Similar challenges have been noted in studies from Bangladesh and Kenya, where unregulated antibiotic use is common due to weak enforcement of regulations (Chowdhury *et al*., 2022; Kariuki *et al*., 2023; Kisoo *et al*., 2023).

Farmers’ reliance on agrovets and chemists for antibiotic procurement, rather than veterinarians or official channels, suggests a lack of proper regulation and monitoring. The low rate of veterinary prescriptions (8.99%) indicates a need for stronger regulation and oversight by authorities such as the Zambia Medicines Regulatory Authority (ZAMRA) and the Veterinary Council of Zambia (VCZ). Similar regulatory challenges are faced in Southeast Asia and parts of South America, where unregulated antibiotic markets contribute to widespread misuse and the acceleration of AMR (Chi *et al*., 2017; Miranda, Godoy and Lee, 2018; Chowdhury *et al*., 2022).

This study underscores the need for comprehensive, multi-faceted interventions to address AMU and AMR in Zambia’s aquaculture sector. Key strategies should include strengthening regulatory frameworks to monitor antibiotic use, enhancing access to veterinary services, and promoting biosecurity practices at the farm level. Additionally, targeted education and training programs are essential to bridge the knowledge gaps among farmers, especially those with less experience or smaller production capacities.

Drawing lessons from countries such as Norway and Scotland, which have successfully reduced antibiotic use through strict regulation, robust disease monitoring, and effective biosecurity measures, Zambia could adopt similar approaches. Moreover, collaboration between government bodies, academic institutions, and the private sector could facilitate the development of a sustainable aquaculture industry that prioritizes responsible AMU and mitigates the risk of AMR.

## 5. Conclusion

The findings from this study highlight significant gaps in farmers’ knowledge, attitudes, and practices regarding AMU and AMR in Zambia’s aquaculture sector. While there is a generally positive attitude towards responsible antibiotic use, actual practices fall short, particularly in terms of adherence to biosecurity protocols and proper antibiotic use. The challenges observed in Zambia are reflective of broader trends in developing countries, where rapid aquaculture expansion has outpaced regulatory and educational efforts. Targeted interventions focusing on farmer education, regulatory enforcement, and improved access to veterinary services are crucial to promoting responsible AMU and curbing the spread of AMR.

## 6. Acknowledgements

We would like to express our sincere gratitude to the fish farmers in the study areas for their cooperation and support as they were crucial for the successful completion of this research. We also extend our heartfelt appreciation to the Food and Agriculture Organization (FAO) Zambia for their financial and logistical support. Additionally, we are grateful for the facilitation provided by the Ministry of Fisheries and Livestock through the Department of Fisheries whose assistance was essential in ensuring the smooth execution of this study.

## 7. Author’s contribution

KN, GM and CM conceptualized and designed the study. KN, MN coordinated data collection. KN performed data analysis, and drafted the manuscript. MC, GM, and JMB provided technical support and contributed to result interpretation. CC, MS1, MS2, and TK provided input and reviewed the manuscript. BMH and NMM supervised the study and reviewed the manuscript. All authors approved the final manuscript.

## 8. Competing Interests

The authors declare that there are no conflicts of interest associated with this study.

## Notes

### Competing Interest Statement

The authors have declared no competing interest.

